# Development and evaluation of novel quantitative PCR (qPCR) and loop-mediated isothermal amplification (LAMP) assays for bovine adenovirus type 7

**DOI:** 10.1101/2025.11.06.687058

**Authors:** Mohamed Kamel, Josiah Levi Davidson, Andres Dextre, Aaron Ault, Deepti Pillai, Jennifer Koziol, Jon P. Schoonmaker, Timothy A. Johnson, Mohit S. Verma

**Affiliations:** Department of Agricultural and Biological Engineering, Purdue University, West Lafayette, IN, USA, 47907; Department of Medicine and Infectious Diseases, Faculty of Veterinary Medicine, Cairo University, Giza 12211, Egypt; Birck Nanotechnology Center, Purdue University, West Lafayette, IN, USA, 47907; Department of Electrical and Computer Engineering, Purdue University, West Lafayette, IN, USA, 47907; Department of Comparative Pathobiology, Purdue University, West Lafayette, IN, USA, 47907; Department of Veterinary Clinical Science, Purdue University, West Lafayette, IN, USA, 47907; School of Veterinary Medicine, Texas Tech University, Amarillo, TX, USA, 79106; Department of Animal Sciences, Purdue University, West Lafayette, IN, USA, 47907; Weldon School of Biomedical Engineering, Purdue University, West Lafayette, IN, USA, 47907

**Keywords:** bovine adenovirus type 7, bovine respiratory disease, loop-mediated isothermal amplification, diagnostics, qLAMP, qPCR, BAV-7, BAdV7

## Abstract

Herein, we report the development of molecular assays for the specific detection of bovine adenovirus type 7 (BAV-7), a prevalent pathogen associated with bovine respiratory disease (BRD). To overcome the limitations of the current diagnostic methods, we developed and optimized a TaqMan quantitative polymerase chain reaction (qPCR) assay with a limit of detection (LOD) of 50 copies per reaction (10 copies/µL) and 100% analytical specificity, showing no cross-reactivity with nine other viral pathogens commonly linked to BRD. Additionally, we designed a fluorescent quantitative loop-mediated isothermal amplification (qLAMP) assay demonstrating equivalent LOD and specificity. Both assays were evaluated using 24 bovine nasal swabs from BRD animals, revealing a 92% agreement between qPCR and qLAMP results. These assays provide powerful tools for advancing our understanding of BAV-7 epidemiology and improving disease management strategies. The qLAMP assay, with its rapid and isothermal amplification, presents potential for future development as a field-deployable diagnostic tool to enable timely detection of BAV-7 in cattle.

## 1. Introduction

Bovine respiratory disease (BRD) is a complex, multifactorial condition affecting cattle worldwide, caused by an interplay of viral and bacterial pathogens (Autio et al., 2007; Centeno- Delphia et al., 2025; Centeno-Martinez et al., 2022; Ellis, 2010, 2009; Fulton, 2009; Goecke et al., 2021; Griffin et al., 2010; Härtel et al., 2004; Jones and Chowdhury, 2010; Kamel et al., 2024; Kishimoto et al., 2017; Klima et al., 2014; Ng et al., 2015; Thanthrige-Don et al., 2018). Among these, bovine adenovirus 7 (BAV-7) has emerged as a significant contributor to respiratory and enteric diseases in cattle (Autio et al., 2007; Härtel et al., 2004). BRD imposes a substantial economic burden on the United States beef and dairy industry, with annual costs exceeding one billion dollars (Edwards, 2010; Griffin, 1997; Hilton, 2014; Kamel et al., 2024; Kirchhoff et al., 2014; McGill and Sacco, 2020; Rice et al., 2007; Smith, 1998; Wilson et al., 2017). This economic impact underscores the critical need for improved diagnostic methods to enhance prevention and control strategies.

BAV-7, one of ten recognized serotypes of Bovine Adenoviruses, belongs to the Atadenovirus genus. It has been detected in multiple countries and contributes significantly to respiratory and enteric diseases in cattle (Cai et al., 1990; Härtel et al., 2004; Hu et al., 1984; Inaba et al., 1973, 1968; Li and Tikoo, 2002; Matsumoto et al., 1969; Matumoto et al., 1970). Recent genomic sequencing has improved our understanding of BAV-7, revealing high sequence identity between US and Japanese isolates (Jesse et al., 2022; Kumagai et al., 2021; Paim et al., 2021). However, current diagnostic techniques for BAV-7, including virus isolation, histopathology, serology, virus neutralizing tests, and conventional polymerase chain reaction (PCR), are limited by factors such as high costs, specialized equipment requirements, potential false negatives, long processing times, complexity, or insufficient specificity (Imus et al., 2019; Inaba et al., 1973; Kishimoto et al., 2017; Wong and Xagoraraki, 2010). These limitations hinder effective detection and control of BAV-7, particularly in field settings or resource-limited areas.

To address these challenges, this study introduces two molecular diagnostic techniques: a real-time TaqMan quantitative PCR (qPCR) assay and a fluorescent quantitative loop-mediated isothermal amplification (qLAMP) assay. The novel real-time TaqMan qPCR assay developed in this study is optimized for BAV-7 detection with a good limit of detection (LOD) and high analytical specificity in laboratory settings. This assay offers precise viral load quantification, which is crucial for disease progression monitoring and research purposes. qPCR has become an indispensable tool in molecular biology and diagnostics due to its ability to detect and quantify specific DNA sequences with high sensitivity and specificity (Heid et al., 1996; Kubista et al., 2006). The TaqMan probe-based qPCR, in particular, offers enhanced specificity through the use of a fluorogenic probe that hybridizes to the target sequence between the forward and reverse primers (Holland et al., 1991; Livak et al., 1995). This technology has been widely applied in various fields, including pathogen detection, gene expression analysis, and genetic testing (Bustin et al., 2009; Espy et al., 2006; Mackay et al., 2002).

Complementing the qPCR assay, we have also developed a fluorescent qLAMP assay, which is the first of its kind specifically designed for BAV-7 detection. LAMP has emerged as a promising tool for point-of-need testing, offering rapid and cost-effective nucleic acid amplification (Ahmed et al., 2025; Notomi et al., 2015, 2000; Obande and Banga Singh, 2020; Parida et al., 2008; Soroka et al., 2021; Wang et al., 2022). LAMP’s ability to amplify DNA at a constant temperature using specific primers and a high-fidelity DNA polymerase makes it ideal for portable diagnostics (Nagamine et al., 2002; Notomi et al., 2000). The technique has been successfully applied to various infectious diseases and environmental analyses globally (Davidson et al., 2021; Goto et al., 2009; Kamel et al., 2025; Mohan et al., 2021; Pascual-Garrigos et al., 2021; Ranjbaran and Verma, 2022; Tomita et al., 2008; Wang et al., 2024b, 2024a, 2023, 2022, 2021).

By developing these two distinct assays, we provide a versatile toolkit that caters to various diagnostic scenarios, from precise laboratory analysis to the potential for rapid field testing. Each technique can be used independently, addressing different needs in BAV-7 detection. The availability of both assays enhances the overall flexibility and effectiveness of BAV-7 diagnosis, ultimately contributing to improved control and prevention strategies for BRD. This approach allows for adaptable diagnostic strategies, addressing the need for both highly accurate detection in laboratory settings and rapid screening in field conditions.

These advancements in diagnostic techniques will significantly contribute to improved control and prevention strategies for BAV-7, potentially reducing the economic impact of BRD on the livestock industry globally. The following sections will detail the development, validation, and potential applications of these novel diagnostic assays for BAV-7 detection.

## 2. Materials and Methods

### 2.1 Viruses, plasmids, and genomic DNA preparation

Synthetic genes for plasmid constructs (pUC57), harboring hexon, proteinase, polymerase, and E1B 55K, were purchased from GenScript (Piscataway, NJ). We utilized various viruses for cross-reactivity tests: Bovine Rhinitis A Virus (BRAV; SD-1 strain, ATCC, catalog number: VR-668), Bovine Rhinitis B Virus (BRBV (EC-11; ATCC, catalog number: VR-1806), Bovine Coronavirus (BCV; Mebus, BEI, catalog number: NR-445), Influenza D Virus (IDV; D/bovine/660, kindly provided by Dr. Benjamin Hause (South Dakota State University, USA)), Bovine Adenovirus 3 (BAV-3; WBR-1 strain, ATCC, catalog number: VR-639), Bovine Herpesvirus type 1 (BHV-1; Los Angeles strain, ATCC, catalog number: VR-188), Bovine Viral Diarrhea Virus type 1 (BVDV-1; NADL strain, ATCC, catalog number: VR-534), Bovine Respiratory Syncytial Virus (BRSV; A 51908 strain, ATCC, catalog number: VR-1339), and Bovine Parainfluenza Virus type 3 (BPI3V; SF-4 strain, BEI, catalog number: NR-3234).

Additionally, the BAV-7 strain Fukuroi VR-768™ was sourced from ATCC ((ATCC VR-768) and propagated in MDBK cells. The MDBK cells were cultured in Dulbecco’s Modified Eagle Medium (DMEM) supplemented with 10% fetal bovine serum and maintained at 37°C in a 5% CO_2_ incubator. When the cell monolayer reached 80-90% confluency, it was washed with calcium- and magnesium-free Dulbecco’s phosphate-buffered saline (DPBS) and inoculated with 500 µL BAV-7. Infected MDBK cells were incubated at 37°C for one week, after which a blind passage was conducted using a 1:2 ratio of infected to uninfected MDBK cells. The passaged cells were monitored daily for signs of cytopathic effects (CPE), and the virus was harvested approximately 8 days post-passage after observing extensive cell detachment and rounding. The virus-containing culture was collected and centrifuged to remove cellular debris. The supernatant was then kept in aliquots at -80 °C for further use. The viral genomic DNA was extracted and purified using the PureLink™ Viral RNA/DNA Mini Kit (12-280-050, Invitrogen/Thermo Fisher Scientific) according to the manufacturer’s protocols.

### 2.2 Sample collection and DNA extraction

The Purdue University Animal Care and Use Committee has approved all animal procedures under protocol #1906001911. Nasal swab samples were collected from antibiotic- treated calves exhibiting BRD (n = 24), which were obtained from Elanco. The sampling followed the approved protocol using sterile rayon-tipped double swabs (BD 220135, Becton Dickinson, Franklin Lakes, NJ, USA) and was done following the same previously reported procedures (Centeno-Martinez et al., 2022; Pascual-Garrigos et al., 2021). Briefly, each calf was secured in a livestock handling chute with a head restraint to limit movement. Excess mucus was removed from the nostrils using paper towels. To collect nasal swab samples, a dual-tipped swab was sequentially inserted approximately 5 cm into each nostril and rotated against the mucosal lining for around 5 seconds per nostril. The swabs were then placed into labeled 15 mL conical tubes and transported on ice to the laboratory for processing. Clinical signs indicative of BRD were assessed using the depression, appetite, respiration, and temperature (DART) system, which evaluates depression, anorexia, abnormal respiratory characteristics, and elevated body temperature (Griffin et al., 2010). Nasal swabs were processed by adding 1 mL of nuclease-free water to the swab tips, mixing for 5 minutes via vortex, removing the swab tips, and centrifuging (6000 × g for 10 minutes) to obtain the supernatant. Viral DNA/RNA was extracted from these supernatants following the manufacturer’s guidelines by the PureLink® Viral RNA/DNA Mini Kit (12-280-050, Invitrogen/Thermo Fisher Scientific). The extracted genomic viral RNA/DNA was kept at -80 °C for subsequent analyses.

### 2.3 Primer design

LAMP primer sets were designed to target the conserved regions within the hexon, polymerase, and proteinase genes. These regions were determined through multiple sequence alignment utilizing Clustal W and designed using the LAMP Primer Explorer v5 program (http://primerexplorer.jp/e). The *in silico* specificity of the primers was confirmed via NCBI BLAST analysis (as explained in the next section on *in silico* cross-reactivity). Each primer set comprised six primers: an outer backward primer (B3), an outer forward primer (F3), a backward inner primer (BIP), a forward inner primer (FIP), a loop backward primer (LB), and a loop forward primer (LF). The inclusion of LB and LF was intended to augment the number of loops in the LAMP reaction, thereby accelerating the reaction speed significantly. The LAMP primer sets are detailed in Tables 1 and Table S1 and were purchased from Invitrogen (Carlsbad, CA) in desalted form.

In parallel, qPCR primers and probes for the BAV-7 strains were designed based on the conserved sequences of the BAV-7 strains available in the NCBI database, employing the PrimerQuest tool from Integrated DNA Technologies (IDT) (Table 1). The primer specificity was upheld through NCBI BLAST verification. The qPCR primers were also sourced from Invitrogen (USA) in desalted form, while a HPLC-purified and double-quenched probe was obtained from IDT (USA).

**Table 1:**
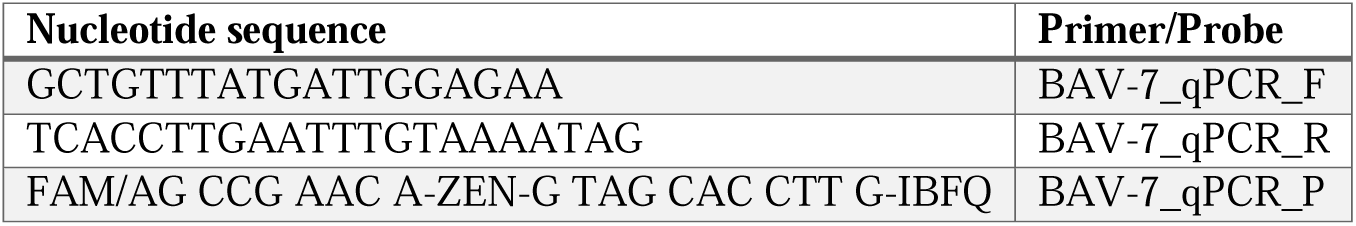
qPCR Primers and Probes Targeting BAV-7 (E1B 55K gene)

### 2.4 In silico cross-reactivity test for BAV-7 qPCR and LAMP Primers against another BAV 4, 5, 6, and 8

To evaluate the specificity of qPCR and LAMP primers designed for BAV-7, an in silico cross-reactivity test was conducted using the BLASTN tool available on the NCBI BLAST website (https://blast.ncbi.nlm.nih.gov/Blast.cgi). The search was conducted against the nucleotide collection (nr/nt) database, which includes a broad collection of nucleotide sequences from GenBank, RefSeq, and other related databases. This comprehensive database was chosen to ensure thorough cross-reactivity testing. In addition, to refine the search and ensure specificity, the search was limited to sequences from BAV-7 (taxid: 10511), BAV-4 (taxid:70333), BAV-5 (taxid:111166), BAV-6 (taxid:111167), and BAV-8 (taxid:120509). This exclusion was necessary to assess the potential cross-reactivity of the primers with closely related adenovirus types.

The search parameters were set to enhance the accuracy of the results. The maximum number of target sequences was set to 5000 to capture a wide range of potential matches. The expected threshold (E-value) was set to 0.01 to filter out non-significant matches, ensuring that only meaningful similarities were considered. The word size was set to 11, appropriate for the length of the primer sequences, and the match/mismatch scores were set at 2 for a match and -3 for a mismatch. Gap costs were configured with an existence cost of 5 and an extension cost of 2. Additionally, low-complexity regions were filtered out.

### 2.5 Quantitative polymerase chain reaction (qPCR) assay

qPCR was conducted using the qTOWER^3^ real-time system (Analytik Jena AG, Jena, Germany) and the Luna Universal Probe qPCR Kit (New England Biolabs, Ipswich, MA, USA). For the reaction mixture, 10 µL of 2X Luna Universal Probe Master Mix, 0.8 µL of 10 µM for each forward and reverse primer, 0.4 µL of 10 µM gene-specific probe, a DNA template, and PCR-grade water were combined to a final volume of 20 µL. The thermal cycling conditions included an initial denaturation at 95°C for 1 minute, followed by 45 cycles of denaturation at 95°C for 15 seconds, annealing at 56°C for 15 seconds, and extension at 60°C for 30 seconds.

Prior to optimization, the protocol stipulated an initial denaturation step at 95°C for 1 minute and 45 cycles of denaturation at 95°C for 15 seconds with an annealing and extension temperature of 60°C for 30 seconds according to the manufacturer’s instructions.

### 2.6 qPCR optimization

Optimization of the experimental conditions was accomplished by employing a gradient annealing temperature range of 53.3°C to 61.6°C, utilizing the real-time qTOWER^3^ system. This approach was used to identify the optimal temperature that would yield the maximum fluorescence intensity coupled with the lowest threshold cycle (Ct) value.

### 2.7 qPCR LOD

In the LOD experiment, a range of synthetic DNA concentrations—500, 250, 125, 50, 25, 5, and down to 1 copy per reaction—were tested alongside an No Template Control (NTC), which consisted of water (instead of the template), using the optimized qPCR assay. Briefly, serial dilutions were prepared such that each 5 µL of the dilution contained the specified number of DNA copies. For each qPCR reaction, 5 µL of the diluted template was added to 15 µL of the remaining reaction mixture.

### 2.8 Fluorescent qLAMP assays

The fluorescent qLAMP assay was conducted using primer sets delineated in Table 2 and S3. The reaction mixture comprised 12.5 µL of WarmStart® LAMP Kit 2X Master Mix E1700 (NEB, Ipswich, USA), 2.5 µL of a 10X concentration LAMP primer mix of 16 μM FIP/BIP, 4 μM LF/LB, 2 μM F3/B3 (yielding final concentrations of 1.6 µM FIP/BIP, 0.4 µM LF/LB, and 0.2 µM F3/B3), and 5 µL of a 10X fluorescent dye (Syto-9, supplied with the NEB LAMP Kit). This 20 µL master mix was dispensed into each well of a 96-well plate. Subsequent to this, 5 µL of purified genomic DNA or molecular grade water for the NTC was added. The plate was then capped, and the assay was incubated at 65 °C.

**Table 2:**
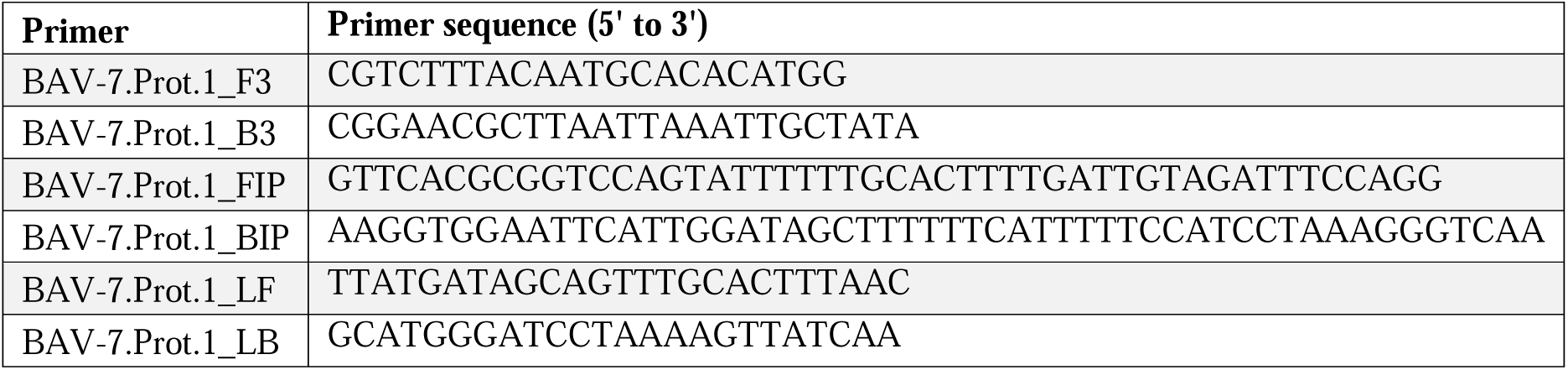
qLAMP BAV-7.proteinase.1 (Prot.1) primer set of the proteinase gene of BAV-7, comprising F2, F1c, B2, B1c ("inner primers"), LB, and LF ("loop primers"), B3 and F3 ("outer primers").

### 2.9 LAMP screening and scoring

Primer sets were evaluated using four replicates of positive control reactions, each containing BAV-7 genomic extract. Various primer sets were tested, with negative control (NTC) reactions using nuclease-free water instead of the BAV-7 genomic extract. Fluorescent qLAMP reactions were carried out as previously detailed in the entitled “Fluorescent LAMP assays”. The assessment criteria (and associated weights), outlined by Mohan *et al* 2021, included average reaction time (20%), standard deviation of reaction time (15%), average maximum fluorescent intensity (5%), standard deviation of maximum fluorescent intensity (5%), and the number of false positives (55%) (Mohan et al., 2021).

False positives were identified as significant increases in fluorescent intensity in NTC reactions, exceeding 10% of the instrument’s maximum intensity. Higher false-positive rates resulted in penalties within the scoring system. Primer sets were ranked according to their total score, with high-scoring ones chosen for further applications. An additional primer set was assigned as a backup in case of issues during subsequent screening. The code for primer scoring can be found in the "Data Availability" section.

### 2.10 Limit of detection experiment for fluorescent qLAMP assay

Ten-fold serial dilutions of synthetic proteinase gDNA of BAV-7 were conducted in nuclease-free water, ranging from 9.5x10^7^ to 1 copy/reaction, with 5 µL of sample used as a template for assessing the LODs. Reactions were performed in triplicate, following the previously described fluorescent LAMP assay protocol. To ascertain the specific LOD of each primer set, the procedure was repeated this time with a narrower range of copy numbers (500, 250, 125, 50, 25, 5, and 1 copies/reaction). The LOD was determined as the lowest concentration with positive amplification in all three replicates.

### 2.11 Spiking experiment for fluorescent qLAMP assay

A synthetic plasmid containing the target proteinase DNA was prepared at concentrations of 500, 250, 125, 50, 25, 5, and 1 copies per reaction. These were then spiked into resuspended nasal swabs collected from BAV-7-negative animals for use in the assay. Two assays were conducted, one with 3 replicates and the other with 6 replicates, and the results were combined for analysis.

### 2.12 Analytical Specificity and cross-reactivity tests for fluorescent qLAMP and qPCR assays

A thorough evaluation of the analytical specificity for fluorescent LAMP and qPCR was performed by testing the genomic extracts from BRAV, BRBV, IDV, BAV-3, BHV-1, BVD-1, BCV, BRSV, and BPI3V. These pathogens are either the common causative agents of BRD, or they induce clinical symptoms in livestock that are similar to those caused by BAV-7. All genomic extract materials were verified using qPCR prior to this analytical specificity analysis.

### 2.13 Evaluating LAMP assay performance using tissue culture-derived viral particles

The frozen supernatant of the tissue culture viral particles, mentioned in the section *Viruses, plasmids and genomic DNA preparation* was spiked in equal volume with resuspended nasal swab samples, and 5 µL was used as a template using the same fluorescent LAMP assay.

### 2.14 Comparative analysis of Fluorescent LAMP and qPCR assays in clinical sample testing

24 nasal swab samples obtained from BRD calves based on a DART diagnosis showing respiratory signs and fever were resuspended in 1 ml water, vortexed, and centrifuged at 6,000 x g for 10 minutes. The resulting supernatant underwent genomic extraction using the PureLink Viral DNA/RNA Mini Kit, adhering to the manufacturer’s protocol. Subsequently, 2 µL of the extracted genomic material served as a template for both fluorescent LAMP and TaqMan probe qPCR analyses.

## 3. Results

### 3.1 Development of qPCR assay: optimization, limit of detection and analytical specificity

Through multiple alignments and analyses of conserved regions within the existing BAV- 7 genomes, we designed primer/probe sequences (Table 1) and confirmed their specificity via a BLAST search against the NCBI database. Our novel optimized qPCR assay for BAV-7 demonstrated good LOD and analytical specificity. Initial attempts using NEB qPCR Universal Probe cycling conditions did not yield satisfactory amplification, fluorescence intensity, or LOD (Fig. S1). We then explored a range of annealing temperatures from 53.3°C to 61.6°C using gradient qPCR to determine the optimal temperature for our assay (Table S1). By optimizing the annealing temperature to 56°C (Fig. 1), we achieved a detection capability of 50 copies per reaction, making our assay suitable for BAV-7 detection (Fig. 2). Additionally, the optimized qPCR assay showed no cross-reactivity with other BRD viruses, confirming its high analytical specificity for BAV-7 (Fig. 3). The *in silico* analysis demonstrated that the qPCR primers (qPCR-F, qPCR-R, and qPCR-P) designed for BAV-7 showed no cross-reactivity with closely related adenoviruses BAV 4, 5, 6, and 8 (Table S2).

### 3.2 Screening of optimal LAMP primers

The genes encoding hexon, polymerase (Poly), and proteinase (Prot) from BAV-7 served as templates for the synthesis of 13 LAMP primer sets (two targeting proteinase, six for polymerase, and five for hexon), as generated by Primer Explorer V5 (Tables 2 and S3). To identify the primer set with the highest efficiency for the fluorescent qLAMP assay, we subjected all primer sets to a uniform evaluation process, which included similar primer concentration, reaction mixture composition, and reaction program parameters, alongside identical template inputs. The primers were assessed for their maximum fluorescence intensity, response time, and specificity in negative controls (Fig. S2). Primer sets were designated as BAV-7.Prot.x or BAV-7.Poly.x or BAV-7.Hexon.x where BAV-7 refers to the target organism, Prot, Poly, Hexon refers to the target gene, and x is an index that increases by one for each new primer set designed. The BAV-7.Prot.1, BAV-7. Poly.2, and BAV-7.Poly.3 primer sets demonstrated the most effective amplification performance among the reliable primers tested, as shown in the scoring result in Table S4. For a more stringent selection, BAV-7.Prot.1 and BAV-7.Poly.2 were chosen for downstream assays. Although BAV-7.Poly.3 demonstrated better scoring results, its NTC at the end of the time threshold was marginally above the baseline and was therefore reserved as a backup along with the other remaining primers.

### 3.3 LOD of the fluorescent qLAMP assay

To ascertain the minimum number of copies that could be detected by the qLAMP method, we conducted an initial series of broad 10-fold serial dilutions using a synthetic DNA template. The preliminary results indicated that the broad LOD of BAV.7.Prot.1 lies within the range of 10 to 100 copies per reaction, as depicted in Fig. S3. Subsequently, we refined the range and tested for the following specific copy numbers per reaction: 500, 250, 125, 50, 25, 5, and 1. The LOD findings, illustrated in Fig. 4, identified an LOD of 50 copies/reaction (or 10 copies/µL of sample). Given this LOD, the fluorescent qLAMP assay established in this study promises to be an effective diagnostic tool for BAV-7 detection.

### 3.4 Analytical specificity of the fluorescent qLAMP assay

The analytical specificity of the LAMP reaction system was subsequently assessed against nine viruses frequently co-occurring with BAV-7 in BRD infections, as depicted in Fig. 5 for BAV-7.Prot.1 and Fig. S4 for BAV-7.Pol.2. Fluorescence indicative of qLAMP was exclusively detected in reactions employing BAV-7 genomic DNA. In contrast, no fluorescence signal was observed in reactions containing genomic extracts of the nine other viruses. These findings indicate the method’s precise ability to identify BAV-7, demonstrating an absence of cross-reactivity with alternative viral pathogens.

The LAMP primers (BAV-7.Prot.1_F3, BAV-7.Prot.1_B3, BAV-7.Prot.1_F1C, BAV- 7.Prot.1_B1C, BAV-7.Prot.1_F2, and BAV-7.Prot.1_B2) exhibited no cross-reactivity with the closely related BAV types 4,5,6, and 8 during *in silico* analysis (Table S5). However, BAV- 7.Prot.1_F2 and BAV-7.Prot.1_B2 showed query coverage of 92% with a 95% percentage identity to two strains of BAV 4 and 6. Additionally, BAV-7.Prot.1_B2 showed similarity to one strain of BAV-4 with 83% query coverage and 95% percentage identity. The low coverage and the fact that only one or two out of eight LAMP primers showed low similarity to one strain of BAV-6 or one strain of BAV-4, respectively, suggest that the overall cross-reactivity should not impact the assay’s analytical specificity.

### 3.5 Contrived samples

Our newly developed fluorescent qLAMP assay successfully detected synthetic BAV-7 DNA spiked in resuspended nasal swab samples (Fig. S5), even in the presence of inhibitors. Notably, it achieved a LOD of 50 copies per reaction in contrived samples (Table 3). This performance extends even beyond synthetic DNA, as the assay also identified viral particles from tissue culture resuspended 1:1 in diluted nasal swab samples in 1 ml water (Fig. S6).

**Table 3:** The limit of detection for fluorescent qLAMP was evaluated using a synthetic BAV-7 proteinase gene plasmid spiked into a resuspended nasal swab matrix. Nine replicate reactions were performed for each concentration. The assay achieved a detection rate of 100% (9/9 replicates) at 500, 250, 125, and 50 copies/ reaction; 77% (7/9) at 25 copies/reaction; 66% (6/9) at 5 copies/reaction; and 55% (5/9) at 1 copy/reaction.

| Copies per Reaction | Positivity (n) | Positivity (%) |
| --- | --- | --- |
| 500 | 9/9 | 100% |
| 250 | 9/9 | 100% |
| 125 | 9/9 | 100% |
| 50 | 9/9 | 100% |
| 25 | 7/9 | 77.78% |
| 5 | 6/9 | 66.67% |
| 1 | 5/9 | 55.56% |

Importantly, this capability eliminates the need for additional sample processing, making it particularly valuable for rapid and accurate BAV-7 detection in clinical settings with limited resources. Overall, these findings highlight the potential significance of our fluorescent qLAMP assay for rapid and precise BAV-7 detection directly from nasal swabs, offering a crucial tool for resource-limited settings.

### 3.6 Evaluation of qPCR and fluorescent qLAMP assays on bovine nasal swabs

To assess the fluorescent qLAMP assay’s efficacy in detecting BAV-7 in clinical specimens, we examined 24 nasal swabs from calves exhibiting signs of BRD. These samples were subjected to nucleic acid extraction and analyzed using both qPCR and fluorescent qLAMP assays. The results are detailed in Table 4. The fluorescent qLAMP identified two samples as positive that qPCR did not detect, while 22 samples were negative by both methods. The high overall agreement of 92% between the fluorescent qLAMP and qPCR assays underscores the fluorescent qLAMP assay’s viability for clinical use. In addition, this study introduces a qPCR assay that specifically targets BAV-7, excluding other adenoviruses, and establishing an effective tool for the diagnosis and monitoring of BAV-7.

**Table 4:** The percentage of overall agreement between the fluorescent qLAMP assay and qPCR assay in assessing viral genomic material extracted with the PureLink® Viral RNA/DNA Mini Kit from 24 calf nasal swab samples.

|  | qPCR |  |  |
| --- | --- | --- | --- |
|  |  | Positive | Negative |
| Fluorescent qLAMP | Positive | 0 | 2 |
|  | Negative | 0 | 22 |
| Overall agreement | 92% |  |  |

## 4. Discussion

Our previous research has focused on bacterial pathogens associated with BRD, specifically *P. multocida*, *M. haemolytica*, and *H. somnus* (Mohan et al., 2021; Pascual-Garrigos et al., 2021). In our current investigation, we are focusing on BAV-7, for which no targeted specific molecular diagnostic test currently exists. BAV-7 is implicated in the complex etiology of BRD, a condition that increasingly compromises cattle production (Hilton, 2014; Mosier, 2014). Traditionally, BAV-7 infections have been confirmed through an array of methods: virus culture, isolation, nonspecific qPCR (BAV 4-8), neutralization tests, serological assays, and PCR for the pan-adenovirus types BAV-4 to BAV-8 (Maluquer de Motes et al., 2004; Sibley et al., 2011; Tsuchiaka et al., 2016). Despite their widespread use, these diagnostic approaches are beset with challenges. PCR may yield false negatives at low viral concentrations, while serological assays can produce false positives due to cross-reactivity with similar viruses. Additionally, the process of virus isolation is not only time-intensive, but also prone to false negatives at low viral loads (Sibley et al., 2011).

Over the last ten years, the speed, sensitivity, and specificity of routine PCR have established it as a pivotal tool for laboratory diagnostics. However, few studies focusing on BAV-7 have employed qPCR assays, which, despite their ability to detect a range of adenovirus serotypes (4-8), are not tailored specifically for BAV-7 (Barnewall et al., 2022; Kishimoto et al., 2017; Tsuchiaka et al., 2016; Wong and Xagoraraki, 2010). With the advent of next-generation sequencing, we gained access to the complete genome sequences of several BAV-7 strains, such as Fukuroi, TS-GT, and SD18-74 (Jesse et al., 2022). Owing to the high sequence identity shared among these strains (exceeding 99%) and their conserved regions, various primer sets targeting the hexon, polymerase, E1B 55K, and proteinase genes were developed for primer design.

qPCR is an essential technique for pathogen detection, including viruses (Fukumoto et al., 2015; López et al., 2003). In virology, crafting specific and sensitive qPCR assays is critical for the precise identification of viruses, thereby facilitating effective disease surveillance and intervention. This study assessed the efficacy of a novel qPCR primer/probe set tailored for the detection of BAV-7 strains. By employing a gradient qPCR method, the optimal annealing temperature was determined to be 56°C. This meticulous optimization yielded an annealing temperature that enhanced the assay’s LOD and analytical specificity.

The findings revealed that, at 56°C, our newly developed qPCR assay achieved a LOD of 50 copies per reaction (10 copies/µL of sample) specifically for BAV-7. This represents an improvement compared to previously reported LODs for BAV detection. Notably, existing qPCR assays that are non-specific to BAV-7 typically employ degenerate primers and probes designed to detect BAV 4-8. One such degenerate qPCR assay reported an LOD of 100 copies per reaction (Kishimoto et al., 2017). Our assay’s enhanced LOD (50 copies per reaction vs. ≤100 copies per reaction) demonstrates its superior performance in detecting low viral loads, particularly when compared to these broader, less specific methods. Moreover, while these qPCR assays typically show LOD of 10 copies per reaction in environmental fecal samples (Wong and Xagoraraki, 2010), our BAV-7-specific assay maintains comparable LOD while offering significantly improved analytical specificity. The new qPCR primer/probe set exhibited high analytical specificity for BAV-7, showing no cross-reactivity with related BRD pathogens or other BAV serotypes. This analytical specificity represents a substantial advancement over the degenerate qPCR approaches, which lack the specificity needed for serotype-specific diagnosis and epidemiological studies.

Our method thus mitigates the risk of false-positive outcomes and significantly improves the accuracy of BAV-7 diagnosis in clinical and research settings. The combination of good LOD and high analytical specificity in our assay addresses critical limitations in existing diagnostic methods, including the degenerate qPCR assays for BAV 4-8. This improvement is particularly valuable for early detection of BAV-7 infection, for monitoring low-level viral presence in carrier animals or environmental samples, and for distinguishing BAV-7 from other BAV serotypes in mixed infections or complex clinical scenarios.

In addition to the qPCR assay, we developed a novel fluorescent qLAMP assay. In our testing, two sets of LAMP primers (polymerase and proteinase) demonstrated high analytical specificity for BAV-7 and effectively excluded viruses associated with other BRD pathogens. The novel fluorescent qLAMP assay’s analytical specificity for BAV-7 was assessed against a panel of nine distinct viruses. This new LAMP assay exhibited a detection threshold of 50 copies per reaction (10 copies/µL). Moreover, it displayed high analytical specificity, showing no cross-reactivity with BRD-related viruses, thus confirming its capability to precisely detect BAV-7 without yielding false positives. Spiking experiments in nasal swabs confirmed a LOD at 50 copies per reaction for the fluorescent qLAMP assay. Notably, these spiking experiments were performed directly on resuspended nasal swab samples without the need for DNA extraction, which represents a significant advantage over the qPCR assay that requires nucleic acid purification. When applied to spiked nasal swabs and crude tissue culture supernatants, these primers reliably identified viral particles, verifying their practical usefulness in actual sample matrices. When comparing the amplification curves for extracted BAV-7 genomic DNA and spiked tissue culture viral particles, a noticeable difference in detection time is observed. The genomic extract produces faster and stronger amplification, while the signal from spiked tissue culture viral particles appears later and with more variability. This difference is likely due to a combination of factors. First, the concentration of viral DNA in purified genomic extracts is typically higher and more consistent than in crude tissue culture preparations, where the actual number of intact viral particles may be lower or less evenly distributed. Second, extraction efficiency can play a significant role: purified extracts provide direct access to viral DNA, whereas viral particles in tissue culture supernatant may not be fully lysed or released, reducing the amount of template available for amplification. Additionally, inhibitors present in crude samples may further delay or reduce amplification efficiency. The extraction-free protocol reduces processing time, cost, and technical complexity, making the LAMP assay more accessible for field applications. Looking ahead, we aim to expand this research to include colorimetric LAMP and paper LAMP applications on farms, further leveraging this extraction-free advantage for point-of-care diagnostics.

When comparing the detection of BAV-7 in BRD nasal swabs using the developed qPCR and fluorescent qLAMP assays, both demonstrated similar effectiveness, with a 92% concordance rate. This indicates that either assay can reliably diagnose BAV-7. However, despite the comparable LOD of 50 copies per reaction, fluorescent qLAMP displayed a marginally higher detection rate for lower copy numbers, a finding underscored by its ability to identify two positive samples that qPCR failed to detect, suggesting fluorescent qLAMP’s slightly superior performance (potentially because it is less likely to be inhibited by inhibitors either in complex matrix or purified extract) (Nwe et al., 2024; Soroka et al., 2021).

Looking ahead, we aim to expand this research to include colorimetric LAMP and paper LAMP applications on farms, further enhancing the accessibility and practicality of BAV-7 detection in field settings. These advancements in diagnostic techniques hold promise for more effective control and prevention strategies for BRD, potentially reducing its economic impact on the livestock industry globally.

## 5. Conclusions

This study has developed two novel assays for BAV-7 detection: qPCR and fluorescent qLAMP. These innovative diagnostic tools offer three advantages: (i) high analytical specificity (100%) by differentiating BAV-7 from nine closely related BRD pathogens, (ii) rapid detection by providing results in about one hour instead of days or weeks, and (iii) potential for surveillance due to their rapid nature and minimal sample processing needed. For the qPCR assay, sample processing requires nasal swab collection, swab resuspension in nuclease-free water, centrifugation (6000 × g for 10 minutes), and viral DNA extraction using the PureLink® Viral RNA/DNA Mini Kit. The fluorescent qLAMP assay requires similar initial processing steps (swab collection, resuspension) but can also be performed directly on resuspended nasal swab samples without the need for viral nucleic acid extraction or purification, as demonstrated in the spiking experiments.

Despite these advancements, there are two limitations that need to be addressed in the future: (i) further testing is needed across various sample types (e.g., feces and tissue samples), and (ii) the assays need to be evaluated in a larger number of clinical samples with higher BAV- 7 prevalence to determine their clinical performance since the current samples did not have any BAV-7 according to the qPCR test.

Looking ahead, several promising research opportunities emerge: (i) development of colorimetric LAMP format for on-site applications (for example, we have demonstrated this possibility for BRD-associated bacteria (Pascual-Garrigos et al., 2021)) and (ii) creation of portable, rapid detection and surveillance methods for resource-limited settings (for example, we have demonstrated that LAMP could be conducted in the back of a car (Wang et al., 2024a)).

## Acknowledgments

The following reagents were obtained through BEI Resources, NIAID, NIH: BCV, Mebus strain, NR-445; and BPI3V, SF-4 strain, NR-3234.

## Declarations and Disclosures

Ethics approval and consent to participate

Sample collection was conducted under the approval of the Institutional Animal Care and Use Committee (IACUC) of Purdue University, protocol number 1906001911.

## Declaration of AI or AI-assisted Technologies in the Writing Process

The authors used Grammarly (https://grammarly.com/) during manuscript preparation to address grammar and language issues. They then reviewed and edited all content themselves and take full responsibility for the final publication.

## Author Contributions

Mohamed Kamel: Conceptualization, Methodology, Validation, Formal Analysis, Investigation, Data Curation, Writing – Original Draft Preparation, and Writing – Review & Editing. Josiah Levi Davidson: Methodology, Software, and Investigation. Andres Dextre: Methodology and Investigation. Aaron Ault: Conceptualization and Resources. Deepti Pillai: Conceptualization and Methodology. Jennifer Koziol: Conceptualization and Methodology. Jon Schoonmaker: Conceptualization and Resources. Timothy A. Johnson: Conceptualization, Methodology, and Resources. Mohit S. Verma: Conceptualization, Methodology, Resources, Writing – Review & Editing, Supervision, Project Administration, Funding Acquisition.

## Data Availability

The raw data supporting the findings of this study are accessible through Mendeley Data, under the following DOI: https://doi.org/10.17632/zkcxkycfc2.1. The primer scoring code utilized in this research is available in a public repository at the Purdue University GitHub organization, located at: https://github.itap.purdue.edu/VermaLab/PrimerScoring.

---

**Fig. 1:** Optimization of qPCR condition temperatures was assessed using gradient qPCR with annealing temperatures ranging from 53.3 °C to 61.6 °C. Gradient qPCR was performed using synthetic BAV-7 E1B 55K gene DNA plasmid at 10^5^ copies per reaction as a template. Annealing temperatures ranged from 53.3 °C to 61.6 °C. Reactions were run with Luna Universal Probe qPCR Master Mix on a qTOWER3 real-time PCR system. Cycling conditions included initial denaturation at 95 °C for 1 minute, followed by 45 cycles of 95 °C for 15 seconds, annealing at gradient temperatures for 15 seconds, and extension at 60 °C for 30 seconds. This optimization identified the temperature yielding the highest fluorescence and lowest cycle threshold values for sensitive and specific detection of BAV-7.

**Fig. 2:** Limit of detection (LOD) testing of qPCR primers/probe using a range of DNA concentrations of the synthetic E1B 55K gene sequence of BAV-7 (500, 250, 125, 50, 25, 5, and 1 copies/reaction). Each 5 µL of the dilution contained the specified number of DNA copies. For each qPCR reaction, 5 µL of the diluted template was added to 15 µL of the remaining reaction mixture. The qPCR reaction was carried out with an initial denaturation step at 95°C for 1 minute, followed by 45 cycles of 95°C for 15 seconds, 56°C for 15 seconds, and 60°C for 30 seconds.

**Fig. 3:** Analytical specificity and cross-reactivity of the Bovine Adenovirus 7 (BAV-7) quantitative PCR (qPCR) assay. The qPCR assay was conducted to assess the analytical specificity and cross-reactivity of the BAV-7 primers and probe against a panel of related bovine respiratory viruses, including Bovine Rhinitis A Virus (BRAV), Bovine Rhinitis B Virus (BRBV), Influenza D virus (IDV), Bovine Adenovirus 3 (BAV-3), Bovine Herpesvirus type 1 (BHV-1), Bovine Viral Diarrhea Virus type 1 (BVD-1), Bovine Coronavirus (BCV), Bovine Respiratory Syncytial Virus (BRSV), and Bovine Parainfluenza Virus type 3 (BPI3V). Viral genomic DNA or RNA was extracted and used at standardized concentrations of 10^5^ copies per reaction. The qPCR assay was performed using Luna Universal Probe qPCR Kit under optimized cycling conditions: initial denaturation at 95°C for 1 minute, followed by 45 cycles of 95°C for 15 seconds, annealing at 56°C for 15 seconds, and extension at 60°C for 30 seconds. No cross-reactivity was observed with non-BAV-7 viruses, demonstrating high assay specificity.

**Fig. 4:** Fine LOD experiment demonstrating the result of the BAV-7 qLAMP assay of proteinase.1 (Prot.1) primer set using the fluorescent qLAMP assay testing different concentrations of the BAV-7 proteinase synthetic DNA (500, 250, 125, 50, 25, 5, and 1 copies/reaction). Each 5 µL of the dilution contained the specified number of DNA copies. For each qLAMP reaction, 5 µL of the diluted template was added to 20 µL of the remaining reaction mixture

**Fig. 5:** Analytical specificity and cross-reactivity of BAV-7 proteinase.1 (Prot.1) LAMP primer set. Fluorescent LAMP assay for testing BAV-7 proteinase.1 LAMP primer set analytical specificity and cross-reactivity. Each 25 µL reaction contained 12.5 µL WarmStart® LAMP Kit 2X Master Mix (New England Biolabs, E1700), 2.5 µL 10X LAMP primer mix (final: 1.6 µM FIP/BIP, 0.4 µM LF/LB, 0.2 µM F3/B3), 5 µL Syto-9 dye, and 5 µL genomic template. Templates were BAV-7 and nine other viruses: Bovine Rhinitis A Virus (BRAV), Bovine Rhinitis B Virus (BRBV), Influenza D Virus (IDV), Bovine Adenovirus 3 (BAV-3), Bovine Herpesvirus type 1 (BHV-1), Bovine Viral Diarrhea Virus type 1 (BVDV-1), Bovine Coronavirus (BCV), Bovine Respiratory Syncytial Virus (BRSV), and Bovine Parainfluenza Virus type 3 (BPI3V). Incubation was at 65°C for 60 minutes. Only BAV-7 produced a positive result, indicating high specificity.

