## Supplementary Information for "Development and evaluation of novel quantitative PCR (qPCR) and loop-mediated isothermal amplification (LAMP) assays for bovine adenovirus type 7"

**for**

**Supplementary figures**

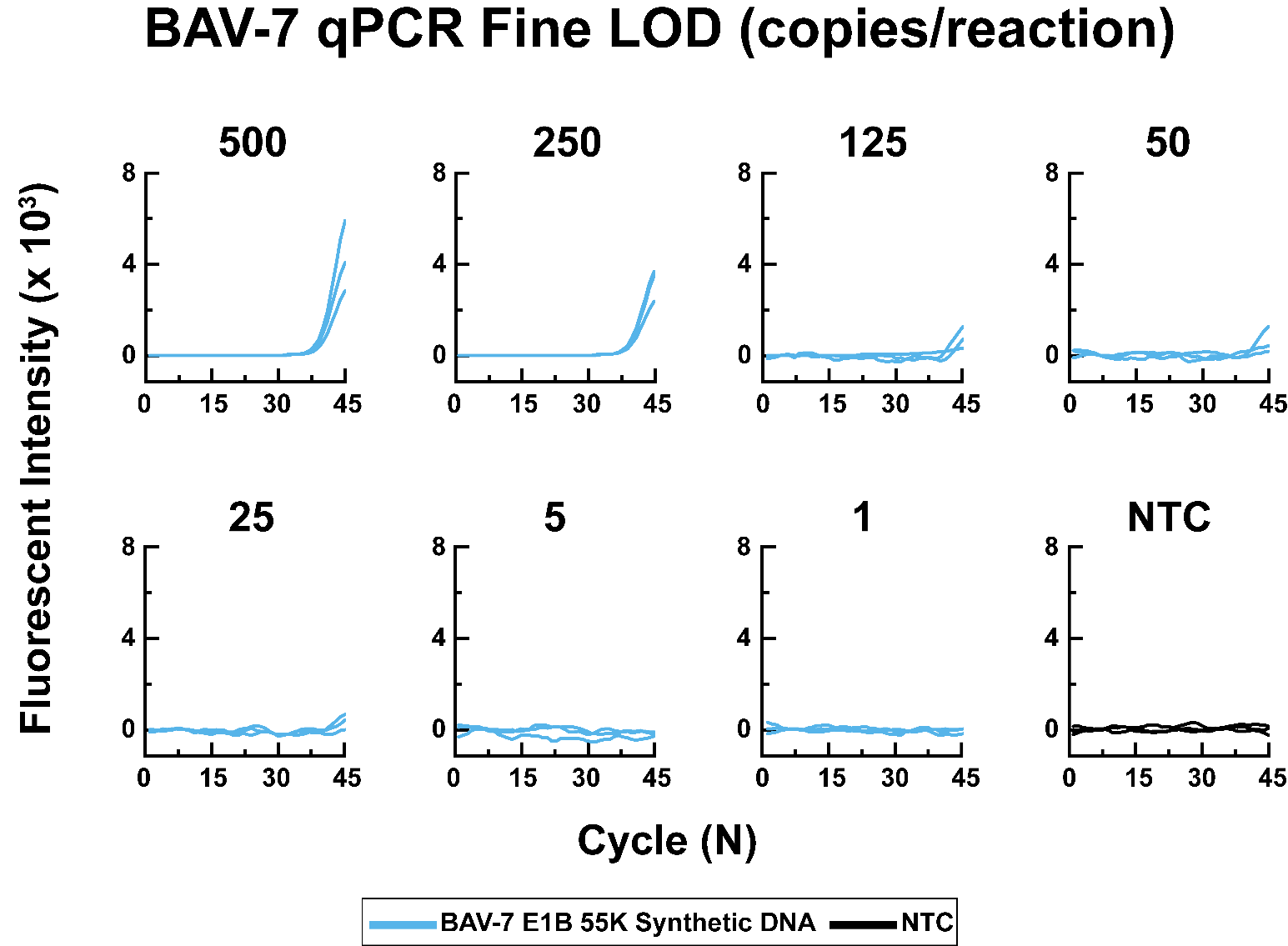

**Fig. S1:** LOD experiment showing the results of the BAV-7 qPCR assay. The assay was performed using the NEB qPCR Universal Probe Kit with different concentrations of 500, 250, 125, 50, 25, 5, and 1 copies per reaction. The qPCR reaction included an initial denaturation step at 95°C for 1 minute, followed by 45 cycles of denaturation at 95°C for 15 seconds, and annealing/extension at 60°C for 30 seconds, according to the manufacturer’s instructions.

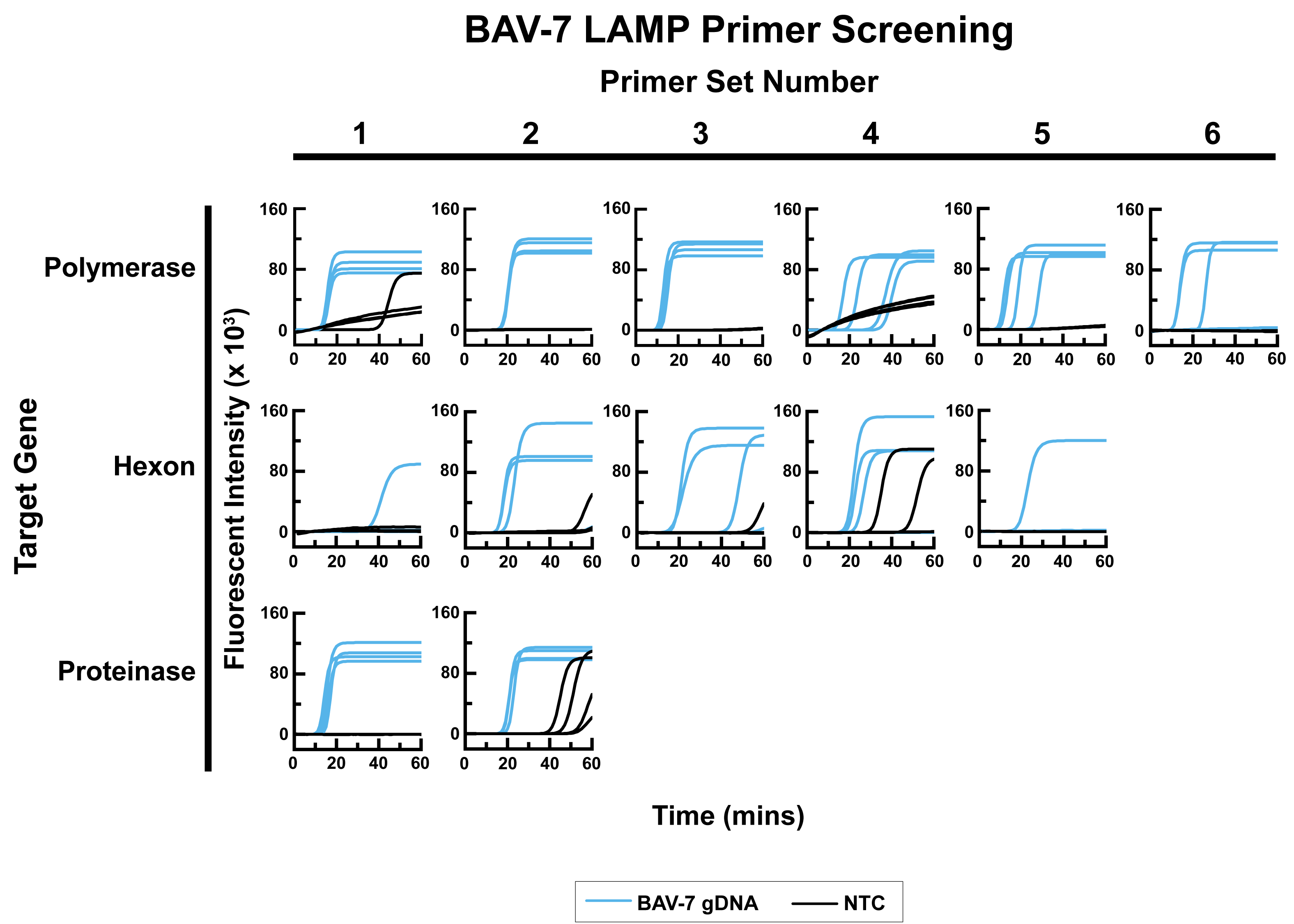

**Fig. S2:** Screening of LAMP primers for BAV-7 hexon (5 primer sets), polymerase (6 primer sets), and proteinase genes (2 primer sets) using fluorescent LAMP assay using NEB warm start DNA/RNA LAMP kit and genomic viral extract as a template. Each reaction (25 µL) contained 12.5 µL WarmStart® DNA/RNA LAMP Kit 2X Master Mix (New England Biolabs, E1700), 2.5 µL 10X LAMP primer mix (final concentrations: 1.6 µM FIP/BIP, 0.4 µM LF/LB, 0.2 µM F3/B3), 5 µL Syto-9 fluorescent dye, and 5 µL purified genomic viral extract as template. Reactions were incubated at 65 °C for 30–60 minutes in a 96-well plate reader.

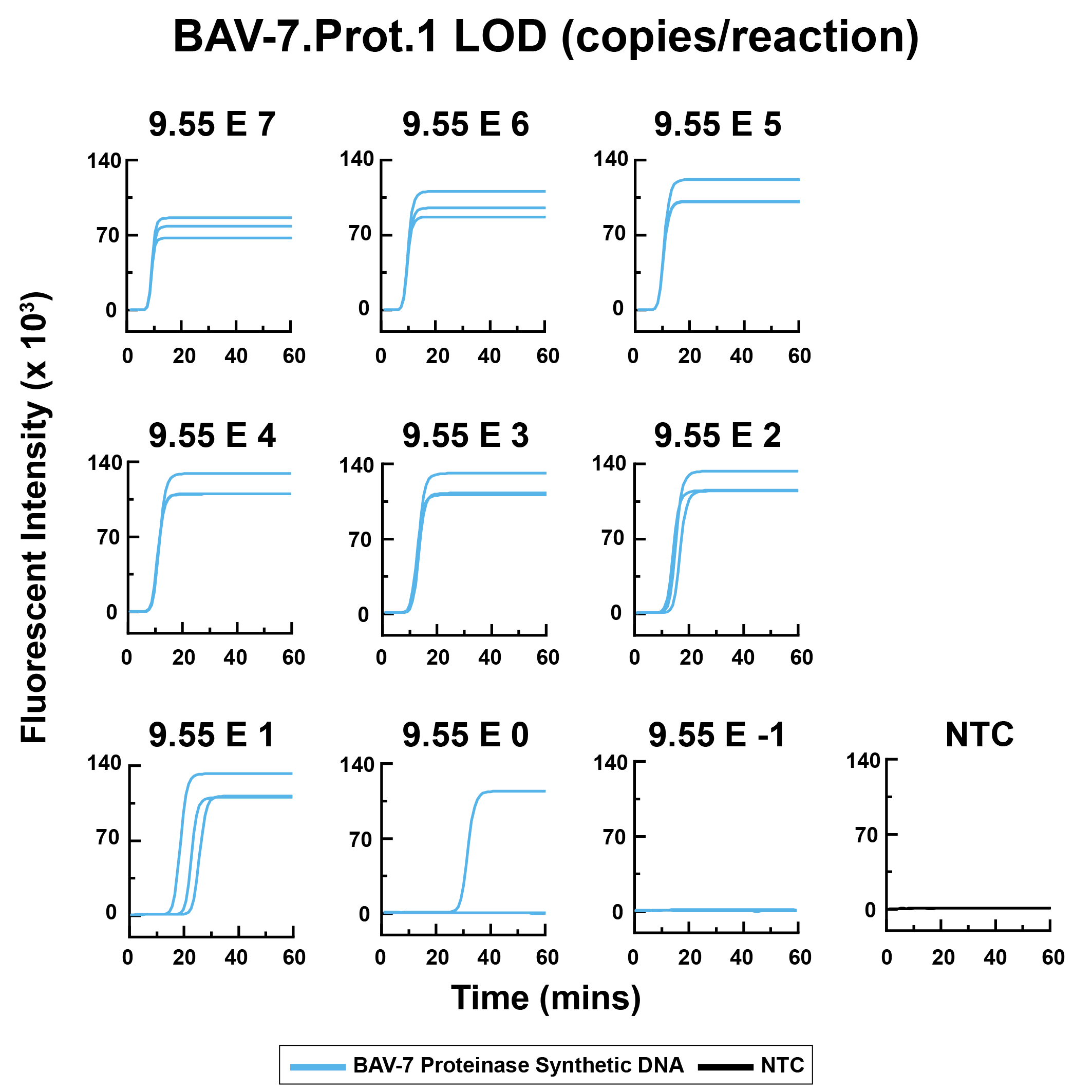

**Fig. S3:** Broad LOD experiment showing the result of the BAV-7 qLAMP assay of proteinase.1 (Prot.1) primer set using the fluorescent qLAMP assay testing a ten-fold serial dilution of a pUC57 plasmid containing the BAV-7 proteinase gene sequence. Ten-fold serial dilutions of pUC57 plasmid encoding the BAV-7 proteinase gene were used as templates in 25 µL reactions containing 12.5 µL WarmStart® LAMP Kit 2X Master Mix (New England Biolabs, E1700), 2.5 µL 10X primer mix (final: 1.6 µM FIP/BIP, 0.4 µM LF/LB, 0.2 µM F3/B3), 5 µL Syto-9 dye, and 5 µL plasmid dilution. Reactions were incubated at 65°C for 60 min.

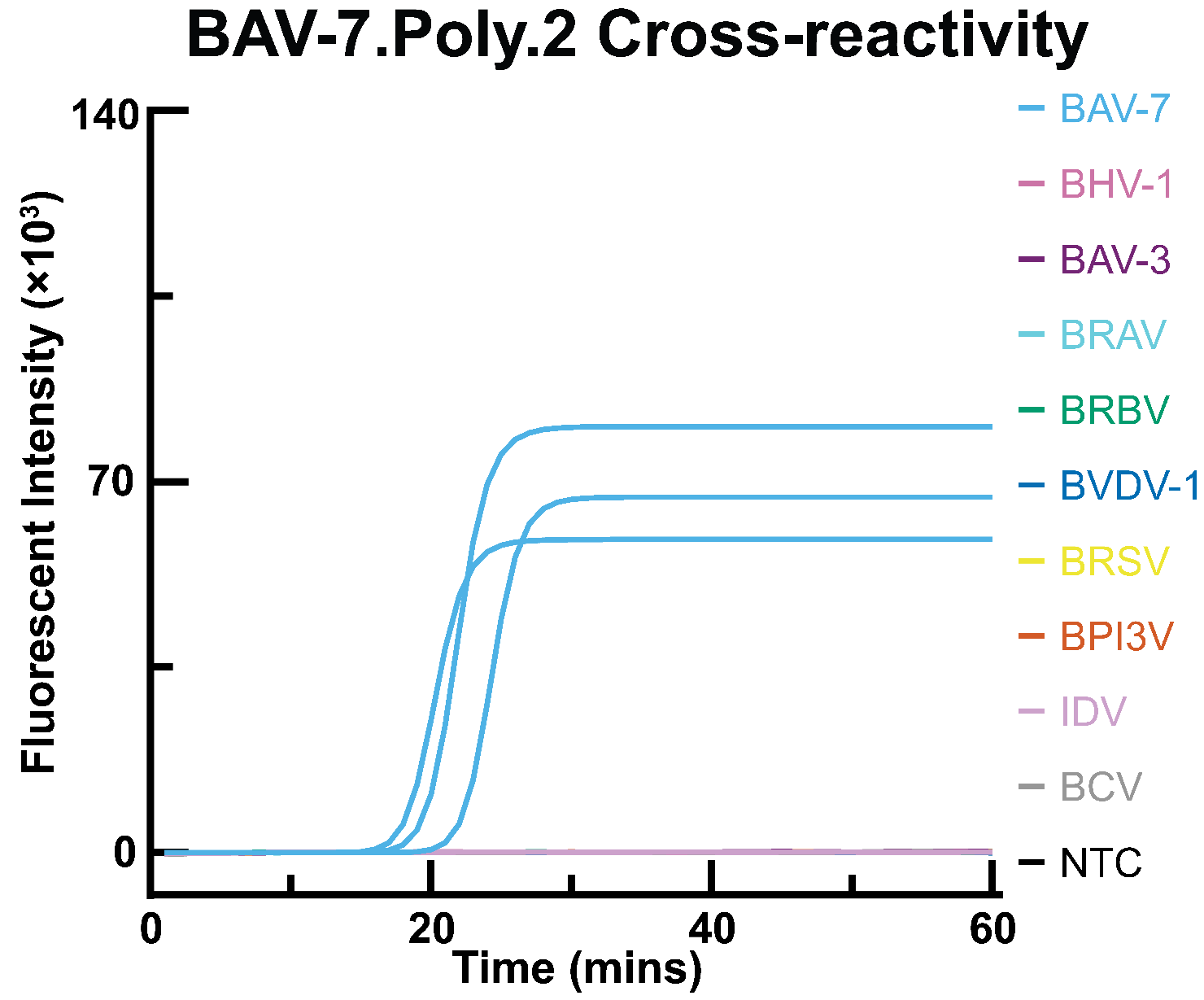

**Fig. S4:** Analytical Specificity and cross-reactivity of BAV-7 polymerase.2 (Pol 2) LAMP primer set. Fluorescent LAMP reactions were prepared in 25 µL volumes using WarmStart® LAMP Kit 2X Master Mix (New England Biolabs, E1700), 2.5 µL of a 10X LAMP primer mix (final concentrations: 1.6 µM FIP/BIP, 0.4 µM LF/LB, 0.2 µM F3/B3), 5 µL Syto-9 fluorescent dye, and 5 µL genomic extract as template. Templates included Bovine Adenovirus type 7 (BAV-7) and genomic extracts from Bovine Rhinitis A Virus (BRAV), Bovine Rhinitis B Virus (BRBV), Influenza D Virus (IDV), Bovine Adenovirus 3 (BAV-3), Bovine Herpesvirus type 1 (BHV-1), Bovine Viral Diarrhea Virus type 1 (BVDV-1), Bovine Coronavirus (BCV), Bovine Respiratory Syncytial Virus (BRSV), and Bovine Parainfluenza Virus type 3 (BPI3V). Incubation was at 65°C for 30–60 minutes. Only reactions containing BAV-7 exhibited an increase in fluorescence, confirming assay specificity.

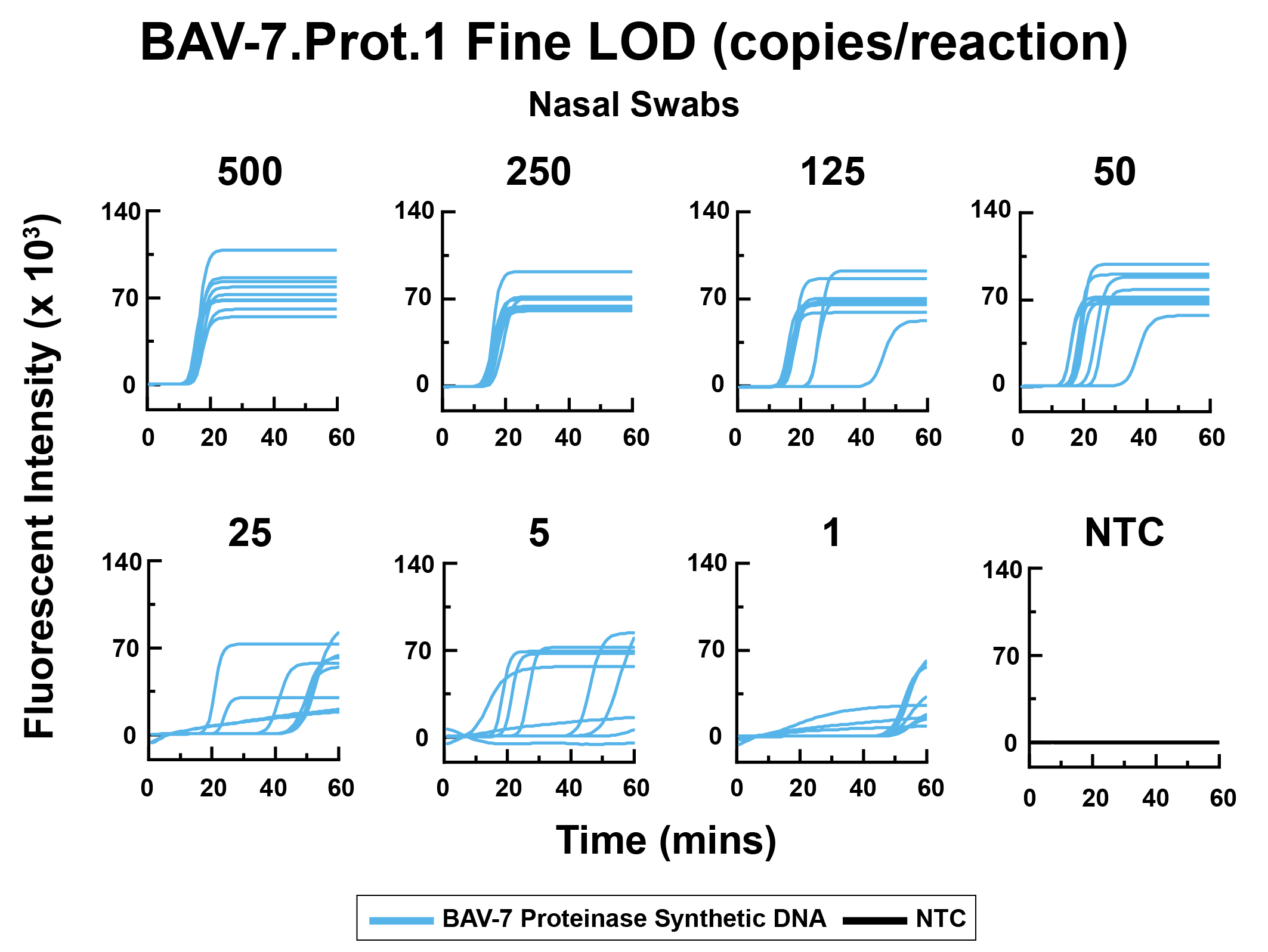

**Fig. S5:** Spiking LOD experiment in resuspended nasal swabs showing the result of the BAV-7 qLAMP assay of proteinase.1 (Prot.1) primer set using the fluorescent qLAMP assay. Each 25 µL fluorescent qLAMP reaction was set up with 12.5 µL WarmStart® LAMP Kit 2X Master Mix (New England Biolabs, E1700), 2.5 µL 10X LAMP primer mix (final concentrations: 1.6 µM FIP/BIP, 0.4 µM LF/LB, 0.2 µM F3/B3), 5 µL Syto-9 fluorescent dye, and 5 µL of resuspended nasal swab matrix spiked with pUC57 plasmid containing the BAV-7 proteinase gene at various concentrations (500, 250, 125, 50, 25, 5, and 1 copies per reaction). Each concentration was tested in 9 technical replicates. Incubation was at 65°C for 60 minutes in a 96-well plate reader.

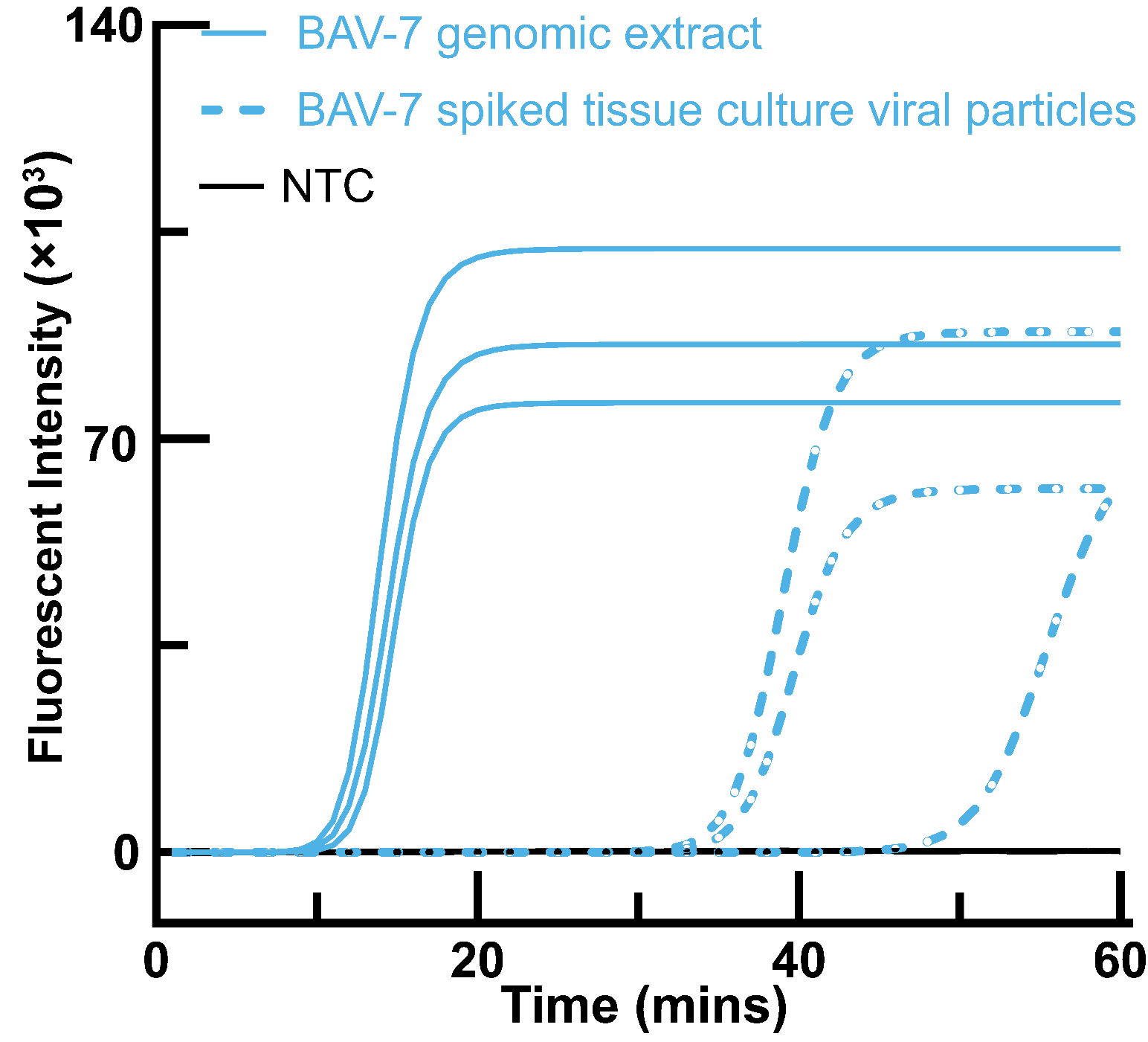

**Fig. S6:** Detection of BAV-7 tissue culture viral particles spiked in nasal swab using qLAMP with genomic extract as a positive control. The viral particles were derived from BAV-7 strain Fukuroi (ATCC VR-768) propagated in MDBK cells cultured in DMEM supplemented with 10% fetal bovine serum at 37°C in 5% CO₂. Virus-containing supernatant was harvested approximately 8 days post-infection after observing cytopathic effects. The exact quantity of viral particles in the supernatant used for spiking is unknown. Five microliters of the spiked sample was used as template in a 25 µL fluorescent qLAMP reaction (12.5 µL WarmStart® LAMP Kit 2X Master Mix [New England Biolabs, E1700], 2.5 µL 10X primer mix at final concentrations of 1.6 µM FIP/BIP, 0.4 µM LF/LB, 0.2 µM F3/B3, and 5 µL Syto-9). A genomic BAV-7 extract was included as a positive control. Incubation was at 65°C for 60 minutes.

**Supplementary tables**

**Table S1:** Assessment of optimal qPCR annealing temperatures: gradient evaluation from 53.3°C to 61.6°C and impacts on Ct values and amplification efficiency. Each 20 µL reaction comprised 10 µL Luna Universal Probe qPCR Master Mix (New England Biolabs), 0.8 µL each of 10 µM forward and reverse primers, 0.4 µL of 10 µM TaqMan probe, 5 10 µL synthetic 10^5^ E1B 55K gene DNA template, and completed to 20 µL with 3 µL PCR-grade water. qPCR was run on a qTOWER3 real-time PCR system with initial denaturation at 95°C for 1 minute, followed by 45 cycles of 95°C for 15 seconds, annealing at gradient temperature for 15 seconds, and 60°C for 30 seconds. The table reports cycle threshold (Ct) values.

| Temperature (°C) | Ct 1 | Ct 2 | Ct 3 | Average Ct Value |
| --- | --- | --- | --- | --- |
| 53.3 | 13.10 | 13.01 | 13.06 | 13.06 |
| 53.5 | 13.13 | 12.82 | 12.93 | 12.96 |
| 54.1 | 12.9 | 12.76 | 12.91 | 12.86 |
| 55 | 12.91 | 12.48 | 12.77 | 12.72 |
| 56 | 12.89 | 12.30 | 12.59 | 12.59 |
| 57 | 13.41 | 13.06 | 13.07 | 13.18 |
| 58 | 13.45 | 13.28 | 13.41 | 13.38 |
| 59 | 13.85 | 13.67 | 13.98 | 13.83 |
| 60 | 14.49 | 13.80 | 13.15 | 13.81 |
| 60.9 | 14.50 | 13.97 | 14.53 | 14.33 |
| 61.6 | 14.34 | 14.24 | 14.65 | 14.41 |

**Table S2:** In Silico Cross-Reactivity of BAV-7 qPCR-F, qPCR-R, qPCR-P Against BAV 4, 5, 6, and 8

This table shows the in-silico analysis results of the qPCR primers designed for BAV-7 to evaluate their cross-reactivity against closely related adenoviruses, including BAV 4, 5, 6, and 8. The analysis was conducted using the NCBI BLASTN tool. Only strains that showed cross-reactivity with the primer sets are included here. Other viruses did not show any cross-reactivity.

| Primer | Description | Scientific Name | Max Score | Total Score | Query Cover | E value | Percentage Identity | Accession Length | Accession |
| --- | --- | --- | --- | --- | --- | --- | --- | --- | --- |
| BAV-7.qPCR_F | Bovine adenovirus 7 strain SD18-74, complete genome | Bovine adenovirus 7 | 35.6 | 35.6 | 100% | 2e-05 | 100.00% | 29973 | OM677816.1 |
|  | Bovine adenovirus 7 strain SD18-74, complete genome | Bovine adenovirus 7 | 35.6 | 35.6 | 100% | 2e-05 | 100.00% | 29933 | MN019492.2 |
|  | Bovine adenovirus 7 TS-GT DNA, complete genome | Bovine adenovirus 7 | 35.6 | 35.6 | 100% | 2e-05 | 100.00% | 30052 | LC605503.1 |
|  | Bovine adenovirus 7 Fukuroi DNA, complete genome | Bovine adenovirus 7 | 35.6 | 35.6 | 100% | 2e-05 | 100.00% | 30034 | LC594788.1 |
| BAV-7.qPCR_R | Bovine adenovirus 7 strain SD18-74, complete genome | Bovine adenovirus 7 | 41.0 | 41.0 | 100% | 8e-07 | 100.00% | 29973 | OM677816.1 |
|  | Bovine adenovirus 7 strain SD18-74, complete genome | Bovine adenovirus 7 | 41.0 | 41.0 | 100% | 8e-07 | 100.00% | 29933 | MN019492.2 |
|  | Bovine adenovirus 7 TS-GT DNA, complete genome | Bovine adenovirus 7 | 41.0 | 41.0 | 100% | 8e-07 | 100.00% | 30052 | LC605503.1 |
|  | Bovine adenovirus 7 Fukuroi DNA, complete genome | Bovine adenovirus 7 | 41.0 | 41.0 | 100% | 8e-07 | 100.00% | 30034 | LC594788.1 |
| BAV-7.qPCR_P | Bovine adenovirus 7 strain SD18-74, complete genome | Bovine adenovirus 7 | 37.4 | 37.4 | 100% | 6e-08 | 100.00% | 29973 | OM677816.1 |
|  | Bovine adenovirus 7 strain SD18-74, complete genome | Bovine adenovirus 7 | 37.4 | 37.4 | 100% | 6e-08 | 100.00% | 29933 | MN019492.2 |
|  | Bovine adenovirus 7 TS-GT DNA, complete genome | Bovine adenovirus 7 | 37.4 | 37.4 | 100% | 6e-08 | 100.00% | 30052 | LC605503.1 |
|  | Bovine adenovirus 7 Fukuroi DNA, complete genome | Bovine adenovirus 7 | 37.4 | 37.4 | 100% | 6e-08 | 100.00% | 30034 | LC594788.1 |

**Table S3:** List of LAMP primer sets screened for hexon (Hex), proteinase (Prot), and polymerase (Pol) genes with primers utilized in the study highlighted in bold

| **Primer** | **Primer sequence (5' to 3')** |
| --- | --- |
| BAV-7.Pol.1_F3 | TCATACGGGCAATAGCCTTT |
| BAV-7.Pol.1_B3 | CGTGCAGGTAAATTGCTTTTTAA |
| BAV-7.Pol.1_FIP | AACTTGGCAAGGATTAAGGTTTATGTTTTTTGATTCGAGTTGGTAAGCAGAA |
| BAV-7.Pol.1_BIP | AAAGAAAGGATACCTTGTTTCCATCTTTTTTGATATAACTTTAGCACTTCCAAAC |
| BAV-7.Pol.1_LF | AACACACTGTTCTTTAAGAGATGC |
| BAV-7.Pol.1_LB | AAACAAGGTTTTTCAAATGCTGG |
| BAV-7.Pol.2_F3 | AGAATCAGTGTCACCATATATACTT |
| BAV-7.Pol.2_B3 | TCCAACAAATCACGTGACATTTAA |
| BAV-7.Pol.2_FIP | ATTTGTACTGGCATGGACTCGTTTTTTGAGTAAGGTTTGCCTCTATCAT |
| BAV-7.Pol.2_BIP | GAAGCTATCTGGGTTGGATACCTTTTTACCTATAACTTATTTAGCAGCAGAT |
| BAV-7.Pol.2_LF | AGTGGGCTGACATATTGTATGG |
| BAV-7.Pol.2_LB | GCGGGTTCGTGATCCATT |
| BAV-7.Pol.3_F3 | GATTTTCTGTCAAGGGTTCTTGC |
| BAV-7.Pol.3_B3 | TGCGAGTCAGAGAAAGAGTTTAA |
| BAV-7.Pol.3_FIP | GACCTTGGAAAGATCAAACTCTGTTTTTTTTGGGTGTTTCTTGTCGTAAG |
| BAV-7.Pol.3_BIP | ATCCGAATTAGTTGAGTTTCTGTCATTTTTCGAACTCTGCAAACAGCAAG |
| BAV-7.Pol.3_LF | TCCAGTGGAAGAACGGATAC |
| BAV-7.Pol.3_LB | GTGAATGGCGCTGTAGTTTT |
| BAV-7.Pol.4_F3 | GTTAGGCAAGGATTCAATTC |
| BAV-7.Pol.4_B3 | GCAGCAGATTGTGATGAT |
| BAV-7.Pol.4_FIP | GGCAAACCTTACTCACAAAGAGGTTCATGTCCTAACTGTGTT |
| BAV-7.Pol.4_BIP | TACAATATGTCAGCCCACTCACCGGTATCCAACCCAGATA |
| BAV-7.Pol.4_LF | ATGGTGACACTGATTCTCTG |
| BAV-7.Pol.4_LB | GAGTCCATGCCAGTACAA |
| BAV-7.Pol.5_F3 | AGTAGGTATCCGTTCTTCC |
| BAV-7.Pol.5_B3 | GCTAAAGGTCATGCCAAA |
| BAV-7.Pol.5_FIP | CGCCATTCACAGTGACAGAGATCCCTTTCCTCAGAGT |
| BAV-7.Pol.5_BIP | CTTGCTGTTTGCAGAGTTCGTAGAACGTCCTACAACCC |
| BAV-7.Pol.5_LF | CGGATATTGAGACCTTGGAA |
| BAV-7.Pol.5_LB | AACTCTTTCTCTGACTCGC |
| BAV-7.Pol.6_F3 | AGAGAATCAGTGTCACCA |
| BAV-7.Pol.6_B3 | GCAGCAGATTGTGATGAT |
| BAV-7.Pol.6_FIP | TCGCGCATTCATGAGTGATGAGTAAGGTTTGCCTCTA |
| BAV-7.Pol.6_BIP | GTCCATGCCAGTACAAATGAAGAAAGAATGGATCACGAACC |
| BAV-7.Pol.6_LF | TGGGCTGACATATTGTATGG |
| BAV-7.Pol.6_LB | CTATCTGGGTTGGATACCG |
| BAV-7.Prot.2_F3 | TTGATTGTAGATTTCCAGGG |
| BAV-7.Prot.2_B3 | GAGATTGGTGGGATTTGG |
| BAV-7.Prot.2_FIP | CCCATGCCAAAGCTATCCAGCTATCATAAATACTGGACCG |
| BAV-7.Prot.2_BIP | TTCCGCTCTATCTTCTCCAGAAAAGACACAGAATAACCCAC |
| BAV-7.Prot.2_LF | GAATTCCACCTTGTTCACG |
| BAV-7.Prot.2_LB | GTACATGTGCCGGATCTT |
| BAV-7.Hex.1_F3 | ACAGATGAAACAAAGGCTTATTTAG |
| BAV-7.Hex.1_B3 | AATAGGTACACGTCCGTTCAT |
| BAV-7.Hex.1_FIP | ATTGGCCCATAAAAACGTTCTTTTTTTGTTACGGAAATATTCCCAGTCTTG |
| BAV-7.Hex.1_BIP | GTAGCTATGTATTTACCTGACAGACTTTTTAGCCATATGAGTTCTGGTCAG |
| BAV-7.Hex.1_LF | TGTAGGTTAGCTGCCAAATTCAT |
| BAV-7.Hex.1_LB | ACTCCAAATGAAGTTATTTTGCCTC |
| BAV-7.Hex.2_F3 | CCTCCAGACAGTGGTAATGAAAT |
| BAV-7.Hex.2_B3 | AGGCAAAATAACTTCATTTGGAGTT |
| BAV-7.Hex.2_FIP | CAAGACTGGGAATATTTCCGTAACTTTTTACTGTAGATGAAGGACGTAACAC |
| BAV-7.Hex.2_BIP | ATGAATTTGGCAGCTAACCTACATTTTTGTCTGTCAGGTAAATACATAGCTAC |
| BAV-7.Hex.2_LF | AAATAAGCCTTTGTTTCATCTGTTT |
| BAV-7.Hex.2_LB | AAGAACGTTTTTATGGGCCAAT |
| BAV-7.Hex.3_F3 | ATGGTGGAGCAGGAAGAA |
| BAV-7.Hex.3_B3 | TGTTGCTTATGGCAGTGG |
| BAV-7.Hex.3_FIP | ATTGCGCCTGTAGCTGCTTTGCAGAGAGTGAAGGAGAA |
| BAV-7.Hex.3_BIP | GGTACCGTCGATAGAGTTAGCGTGACAATCTGGATGAAGTCTG |
| BAV-7.Hex.3_LF | GTGGACGACAATAGGAACCATA |
| BAV-7.Hex.3_LB | GCGTAGTGGCAGTTGACA |
| BAV-7.Hex.4_F3 | CAACCCTAATCCTCAAGTAGG |
| BAV-7.Hex.4_B3 | CTGGATGAAGTCTGTTCACTT |
| BAV-7.Hex.4_FIP | CTGCTTGTGGACGACAATAGGAGGTGGAGCAGGAAGAATC |
| BAV-7.Hex.4_BIP | CAGGCGCAATTTCTACAGAAGCACTACGCCGCTAACTCTAT |
| BAV-7.Hex.4_LF | CTCCTTCACTCTCTGCACTAAG |
| BAV-7.Hex.4_LB | TTGAATTCTACTGGTACCGTCG |
| BAV-7.Hex.5_F3 | TCCTGCTTATGGTTCCTAT |
| BAV-7.Hex.5_B3 | GTTCCAGCATTAGAACCAT |
| BAV-7.Hex.5_FIP | CACTACGCCGCTAACTCTATCTTCTACAGAAGCTATCACCA |
| BAV-7.Hex.5_BIP | ACAGACTTCATCCAGATTGTCATGTAATTCGGTCTGTTGC |
| BAV-7.Hex.5_LF | CGGTACCAGTAGAATTCAAGTA |
| BAV-7.Hex.5_LB | GGAATACACGGATGAAGGAA |

**Table S4:** Analysis of LAMP primer sets designed in this study. Higher overall score is better.

|  |  | Amplification Intensity | | Reaction Time | | False Positive Points Awarded | | | | |  |
| --- | --- | --- | --- | --- | --- | --- | --- | --- | --- | --- | --- |
| Primer Set | **True Positives** | **Average** | **Std. Dev.** | **Avg** | **Std Dev.** | **Total False Positives** | **False Positive 1** | **False Positive 2** | **False Positive 3** | **False Positive 4** | **Overall Score** |
| BAV-7.Poly.3 | 4 | 108914.41 | 8311.54 | 10.75 | 0.83 | 0 | 6 | 12 | 18 | 24 | 97.22 |
| BAV-7.Prot.1 | 4 | 107019.03 | 10550.34 | 12.50 | 1.12 | 0 | 6 | 12 | 18 | 24 | 92.40 |
| BAV-7.Poly.2 | 4 | 110588.64 | 8915.87 | 17.75 | 0.43 | 0 | 6 | 12 | 18 | 24 | 88.22 |
| BAV-7.Poly.5 | 4 | 102284.43 | 6623.98 | 15.50 | 6.58 | 0 | 6 | 12 | 18 | 24 | 84.13 |
| BAV-7.Prot.2 | 4 | 105345.93 | 7856.01 | 18.50 | 0.87 | 4 | 1.59 | 5.31 | 15.39 | 22.49 | 71.21 |
| BAV-7.Poly.1 | 4 | 86954.07 | 12204.02 | 12.75 | 0.43 | 4 | 2.05 | 5.29 | 6.35 | 8.25 | 49.27 |
| BAV-7.Poly.4 | 4 | 97994.93 | 5777.29 | 25.75 | 9.07 | 4 | 0 | 0 | 0 | 0 | 7.34 |
| BAV-7.Hexon.1 | 1 | 0 | 0 | 0 | 0 | 0 | 0 | 0 | 0 | 0 | 0 |
| BAV-7.Hexon.2 | 3 | 0 | 0 | 0 | 0 | 0 | 0 | 0 | 0 | 0 | 0 |
| BAV-7.Hexon.3 | 3 | 0 | 0 | 0 | 0 | 0 | 0 | 0 | 0 | 0 | 0 |
| BAV-7.Hexon.4 | 3 | 0 | 0 | 0 | 0 | 0 | 0 | 0 | 0 | 0 | 0 |
| BAV-7.Hexon.5 | 1 | 0 | 0 | 0 | 0 | 0 | 0 | 0 | 0 | 0 | 0 |
| BAV-7.Poly.6 | 3 | 0 | 0 | 0 | 0 | 0 | 0 | 0 | 0 | 0 | 0 |

**Table S5:** In Silico Cross-Reactivity of BAV-7 LAMP Primers Against BAV 4, 5, 6, and 8

This table presents the in silico cross-reactivity analysis of LAMP primers designed for BAV-7. The primers analyzed include BAV-7.Prot.1_F3, BAV-7.Prot.1_B3, BAV-7.Prot.1_F1C, BAV-7.Prot.1_B1C, BAV-7.Prot.1_F2, and BAV-7.Prot.1_B2. The NCBI BLASTN tool was used to assess potential cross-reactivity with closely related BAV 4, 5, 6, and 8. Only strains that showed cross-reactivity with the primer sets are included here. Other viruses did not show any cross-reactivity.

| Primers | Description | Scientific Name | Max Score | Total Score | Query Cover | E value | Percentage Ident | Acession. Length | Accession |
| --- | --- | --- | --- | --- | --- | --- | --- | --- | --- |
| BAV-7.Prot.1_F3 | Bovine adenovirus 7 strain SD18-74, complete genome | Bovine adenovirus 7 | 41.0 | 41.0 | 100% | 8e-07 | 100.00% | 30052 | LC606503.1 |
|  | Bovine adenovirus 7 TS-GT DNA, complete genome | Bovine adenovirus 7 | 41.0 | 41.0 | 100% | 8e-07 | 100.00% | 30034 | LC597488.1 |
|  | Bovine adenovirus type 7 gene for proteinase | Bovine adenovirus 7 | 41.0 | 41.0 | 100% | 8e-07 | 100.00% | 900 | X53989.1 |
|  | Bovine adenovirus 7 strain SD18-74, complete genome | Bovine adenovirus 7 | 36.5 | 36.5 | 100% | 3e-05 | 95.45% | 29973 | OM677816.1 |
|  | Bovine adenovirus 7 strain SD18-74, complete genome | Bovine adenovirus 7 | 36.5 | 36.5 | 100% | 3e-05 | 95.45% | 30043 | MN901942.2 |
| BAV-7.Prot.1_B3 | Bovine adenovirus 7 strain SD18-74, complete genome | Bovine adenovirus 7 | 44.6 | 44.6 | 100% | 3e-08 | 100.00% | 29973 | OM677816.1 |
|  | Bovine adenovirus 7 strain SD18-74, complete genome | Bovine adenovirus 7 | 44.6 | 44.6 | 100% | 3e-08 | 100.00% | 30043 | MN901942.2 |
|  | Bovine adenovirus 7 TS-GT DNA, complete genome | Bovine adenovirus 7 | 44.6 | 44.6 | 100% | 3e-08 | 100.00% | 30052 | LC606503.1 |
|  | Bovine adenovirus 7 Fukuroi DNA, complete genome | Bovine adenovirus 7 | 44.6 | 44.6 | 100% | 3e-08 | 100.00% | 30034 | LC597488.1 |
|  | Bovine adenovirus 7 PrBovF protease gene, partial cds | Bovine adenovirus 7 | 44.6 | 44.6 | 100% | 3e-08 | 100.00% | 439 | AY288821.1 |
|  | Bovine adenovirus type 7 gene for proteinase | Bovine adenovirus 7 | 44.6 | 44.6 | 100% | 3e-08 | 100.00% | 900 | X53989.1 |
| BAV-7.Prot.1_LF | Bovine adenovirus 7 strain SD18-74, complete genome | Bovine adenovirus 7 | 46.4 | 46.4 | 100% | 3e-08 | 100.00% | 29973 | OM677816.1 |
|  | Bovine adenovirus 7 strain SD18-74, complete genome | Bovine adenovirus 7 | 46.4 | 46.4 | 100% | 3e-08 | 100.00% | 30043 | MN901942.2 |
|  | Bovine adenovirus 7 TS-GT DNA, complete genome | Bovine adenovirus 7 | 46.4 | 46.4 | 100% | 3e-08 | 100.00% | 30052 | LC606503.1 |
|  | Bovine adenovirus 7 Fukuroi DNA, complete genome | Bovine adenovirus 7 | 46.4 | 46.4 | 100% | 3e-08 | 100.00% | 30034 | LC597488.1 |
|  | Bovine adenovirus type 7 gene for proteinase | Bovine adenovirus 7 | 46.4 | 46.4 | 100% | 3e-08 | 100.00% | 900 | X53989.1 |
| BAV-7.Prot.1_LB | Bovine adenovirus 7 strain SD18-74, complete genome | Bovine adenovirus 7 | 44.6 | 44.6 | 100% | 8e-08 | 100.00% | 29973 | OM677816.1 |
|  | Bovine adenovirus 7 TS-GT DNA, complete genome | Bovine adenovirus 7 | 44.6 | 44.6 | 100% | 8e-08 | 100.00% | 30043 | LC606503.1 |
|  | Bovine adenovirus 7 Fukuroi DNA, complete genome | Bovine adenovirus 7 | 44.6 | 44.6 | 100% | 8e-08 | 100.00% | 30052 | LC597488.1 |
|  | Bovine adenovirus 7 PrBovF protease gene, partial cds | Bovine adenovirus 7 | 44.6 | 44.6 | 100% | 8e-08 | 100.00% | 439 | AY288821.1 |
|  | Bovine adenovirus type 7 gene for proteinase | Bovine adenovirus 7 | 44.6 | 44.6 | 100% | 8e-08 | 100.00% | 900 | X53989.1 |
| BAV-7.Prot.1_F1C | Bovine adenovirus 7 strain SD18-74, complete genome | Bovine adenovirus 7 | 31.9 | 31.9 | 100.00% | 2e-04 | 100.00% | 29973 | OM677816.1 |
|  | Bovine adenovirus 7 strain SD18-74, complete genome | Bovine adenovirus 7 | 31.9 | 31.9 | 100.00% | 2e-04 | 100.00% | 30043 | MN901942.2 |
|  | Bovine adenovirus 7 TS-GT DNA, complete genome | Bovine adenovirus 7 | 31.9 | 31.9 | 100.00% | 2e-04 | 100.00% | 30052 | LC606503 |
|  | Bovine adenovirus 7 Fukuroi DNA, complete genome | Bovine adenovirus 7 | 31.9 | 31.9 | 100.00% | 2e-04 | 100.00% | 30034 | LC597488 |
|  | Bovine adenovirus 7 PrBovF protease gene, partial cds | Bovine adenovirus 7 | 31.9 | 31.9 | 100.00% | 2e-04 | 100.00% | 439 | AY288821.1 |
|  | Bovine adenovirus type 7 gene for proteinase | Bovine adenovirus 7 | 31.9 | 31.9 | 100.00% | 2e-04 | 100.00% | 900 | X53989 |
| BAV-7.Prot.1_B1C | Bovine adenovirus 7 strain SD18-74, complete genome | Bovine adenovirus 7 | 41.0 | 41.0 | 100% | 8e-07 | 100.00% | 29973 | OM677816.1 |
|  | Bovine adenovirus 7 strain SD18-74, complete genome | Bovine adenovirus 7 | 41.0 | 41.0 | 100% | 8e-07 | 100.00% | 30043 | MN901942.2 |
|  | Bovine adenovirus 7 TS-GT DNA, complete genome | Bovine adenovirus 7 | 41.0 | 41.0 | 100% | 8e-07 | 100.00% | 30052 | LC606503.1 |
|  | Bovine adenovirus 7 Fukuroi DNA, complete genome | Bovine adenovirus 7 | 41.0 | 41.0 | 100% | 8e-07 | 100.00% | 30034 | LC597488.1 |
|  | Bovine adenovirus 7 PrBovF protease gene, partial cds | Bovine adenovirus 7 | 41.0 | 41.0 | 100% | 8e-07 | 100.00% | 439 | AY288821.1 |
|  | Bovine adenovirus type 7 gene for proteinase | Bovine adenovirus 7 | 41.0 | 41.0 | 100% | 8e-07 | 100.00% | 900 | X53989.1 |
| BAV-7.Prot.1_F2 | Bovine adenovirus 7 strain SD18-74, complete genome | Bovine adenovirus 7 | 46.4 | 46.4 | 100% | 3e-08 | 100.00% | 29973 | OM6778161 |
|  | Bovine adenovirus 7 strain SD18-74, complete genome | Bovine adenovirus 7 | 46.4 | 46.4 | 100% | 3e-08 | 100.00% | 30043 | MN901942.2 |
|  | Bovine adenovirus 7 TS-GT DNA, complete genome | Bovine adenovirus 7 | 46.4 | 46.4 | 100% | 3e-08 | 100.00% | 30052 | LC606503.1 |
|  | Bovine adenovirus 7 Fukuroi DNA, complete genome | Bovine adenovirus 7 | 46.4 | 46.4 | 100% | 3e-08 | 100.00% | 30034 | LC597488.1 |
|  | Bovine adenovirus type 7 gene for proteinase | Bovine adenovirus 7 | 46.4 | 46.4 | 100% | 3e-08 | 100.00% | 900 | X53989.1 |
|  | Bovine adenovirus 6 strain 671130 complete genome | Bovine adenovirus 6 | 38.3 | 38.3 | 92% | 1e-05 | 95.65% | 30024 | NC020074.1 |
|  | Bovine adenovirus 6 strain 671130 complete genome | Bovine adenovirus 6 | 38.3 | 38.3 | 92% | 1e-05 | 95.65% | 30024 | JQ345700.1 |
|  | Bovine adenovirus 4 strain THT/62, complete genome | Bovine adenovirus 4 | 38.3 | 38.3 | 92% | 1e-05 | 95.65% | 31301 | AF036092.3 |
| BAV-7.Prot.1_F2 | Bovine adenovirus 7 strain SD18-74, complete genome | Bovine adenovirus 7 | 44.6 | 44.6 | 100% | 8e-08 | 100.00% | 29973 | OM677816.1 |
|  | Bovine adenovirus 7 strain SD18-74, complete genome | Bovine adenovirus 7 | 44.6 | 44.6 | 100% | 8e-08 | 100.00% | 30043 | MN901942.2 |
|  | Bovine adenovirus 7 TS-GT DNA, complete genome | Bovine adenovirus 7 | 44.6 | 44.6 | 100% | 8e-08 | 100.00% | 30052 | LC606503.1 |
|  | Bovine adenovirus 7 Fukuroi DNA, complete genome | Bovine adenovirus 7 | 44.6 | 44.6 | 100% | 8e-08 | 100.00% | 30034 | LC5974881 |
|  | Bovine adenovirus 7 PrBovF protease gene, partial cds | Bovine adenovirus 7 | 44.6 | 44.6 | 100% | 8e-08 | 100.00% | 439 | AY288821.1 |
|  | Bovine adenovirus type 7 gene for proteinase | Bovine adenovirus 7 | 44.6 | 44.6 | 100% | 8e-08 | 100.00% | 900 | X53989.1 |
|  | Bovine adenovirus 4 PrBovB protease gene, partial cds | Bovine adenovirus 4 | 32.8 | 32.8 | 83% | 5e-04 | 95.00% | 438 | AY288819.1 |
|  | Bovine adenovirus 4 strain THT/62, complete genome | Bovine adenovirus 4 | 32.8 | 32.8 | 83% | 5e-04 | 95.00% | 31301 | AF036092.3 |
